# Breeding success of Eleonora’s falcon (*Falco eleonorae*) in Cyprus revisited using survey techniques for cliff-nesting species

**DOI:** 10.1101/2020.05.04.077248

**Authors:** Thomas G. Hadjikyriakou, Nikolaos Kassinis, Dimitrios Skarlatos, Pantelis Charilaou, Alexander N. G. Kirschel

## Abstract

The global breeding population of Eleonora’s falcon is distributed from the Canary Islands in the west, across the Mediterranean Sea, to Cyprus in the east. The remoteness of nesting colonies, which are predominantly located on sea cliffs and islets, renders breeding success estimation a challenging task, requiring a composite approach to assess each of the breeding stages. Early estimates of the breeding success of Eleonora’s falcon suggested that Akrotiri colony in Cyprus had the lowest breeding success among all the colonies throughout the species’ breeding range, at a level seemingly unsustainable, suggesting the colony might have been in danger of gradual extinction. Here we use a diversity of survey methods using boat, ground and aerial surveys, with the incorporation of photography and photogrammetry, to reassess the breeding success and the effect of nest characteristics on the Eleonora’s falcon breeding population in Cyprus. During a six-year study, we found that Cyprus hosts ~138 ± 8 breeding pairs and that breeding success equals 1.54 ± 0.85 fledglings per breeding pair, thus considerably higher than previous estimates. In addition, by analyzing temporal variation in breeding and nest characteristics, we found that early breeding and reuse of nests positively influence breeding success, but physical nest characteristics have a limited effect on colony productivity. The range of survey methods employed, as well as the array of photography techniques utilized, highly enhanced the efficiency and accuracy of this study, allowing us to overcome the challenge of inaccessibility of nesting cliffs.

## INTRODUCTION

The plight of raptors remains a major conservation concern, despite population recoveries in many species following global declines caused by organochloride pesticides in the 1950-1960’s (Newton et al 2016). Biocides, lead poisoning, direct persecution, egg collection, electrocutions and collisions on power infrastructure and falconry, continue to threaten raptor populations (del Hoyo et al. 1994; Newton et al 2016). However, the major concern nowadays is habitat degradation, fragmentation and loss (del Hoyo et al. 1994). Long-distance migrating birds are exposed to threats along several parts of their migratory range and thus might be particularly prone to decline compared to sedentary species (Berthold 2001; Newton 2004; Sanderson et al. 2006).

Breeding success is a parameter of species demography (others being mortality, emigration and immigration) of critical importance for raptor status assessment (Steenhof and Newton 2007). Thus, data on breeding success can inform conservation measures (Green 2004). For a reliable estimate of breeding success, surveys should include all the stages of breeding phenology including territory occupancy, clutch size and number of offspring hatched and fledged (Brown 1974; Green 2004). However, when conditions impede such extensive fieldwork, surveys should focus at a minimum on quantifying the number of nestlings that reach fledging age (Steenhof and Newton 2007).

Many raptor species nest in areas that are challenging to access (Newton et al 2016), especially for mammalian predators. As a result, cliff nesting species – particularly sea-cliff nesters - are challenging to approach for data collection, and special techniques need to be used (Green 2004). Combining different survey methods (ground, aerial and boat-based) can improve survey efficiency (Andersen 2007; Steenhoff and Newton 2007). Estimating the stage of nesting (e.g. date that egg was laid or fledgling hatched) would require intensive monitoring of each nest or access to the nest (Green 2004), which is impractical on sea-cliff nesting raptors. Alternatively, footage obtained from a few camera traps covering the whole breeding season duration (e.g., Hadjikyriakou and Kirschel 2016) can be useful in establishing a baseline of nestling development. Using such footage, nestling age estimation of many nests can be accomplished using nestling photographs taken from a distance (Clark 2007).

Nest sites are a limiting factor for raptor populations (Danchin and Wagner 1997; Newton 2010). Even for social raptors, nest sites are limited by topography and predation / disturbance potential, and determined by the existence of suitable nesting positions, such as ledges, holes and crevices (Walter 1979). Also, specific ranges of topographic factors such as nest elevation, aspect and cliff inclination have been found to be preferred by cliff-nesting raptors (Urios and Martínez-Abraín 2006), influencing productivity (Touati et al. 2017). To that end, Digital Elevation Models can enhance species-habitat associations (Urios and Martínez-Abraín 2006; Kassara et al 2011). For insect feeding raptors, precipitation and its expected effect on insect abundance (Wolda 1978; Wolda 1988) could affect clutch size (Ristow 2004; Steenhof and Newton 2007), with well-fed females producing more eggs (Wink et al. 1980). in Cyprus was previously considered in danger of decline, with breeding success estimates of 0.8 – 1.0 fledglings per breeding pair (Walter and Foers 1980). Such low breeding success levels might lead to population decline (Ristow and Wink 1985; Gschweng et al. 2011), because at least 1.2 fledglings per breeding pair has been estimated as a threshold to ensure colony sustainability (Ristow and Wink 1985). Estimates in Cyprus compared poorly with the 2.6 fledglings per breeding pair in Morocco (Walter 1979), which suggested an eastward decreasing trend in breeding success of Eleonora’s falcon (Xirouchakis et al. 2012). Also the bird migration pulse over breeding areas in Cyprus restricts feeding opportunities time-wise to a relatively small dawn to early morning window, further increasing the concerns of decline (Walter 1979). In addition, the species has been found to exhibit strong site tenacity (Ristow 1999), which may create population specific pressures, as isolated colonies such as the ones in Cyprus might have restricted gene exchange with other colonies (Swatschek et al. 1993) as well as low immigration levels (Wink et al. 1987; Gschweng et al. 2011). Eleonora’s falcon is included in Annex I of the EU Directive 2009/147/EC, which requires special conservation measures to ensure its survival and reproduction (European Parliament 2009). It is protected through legislation in the Republic of Cyprus (N.152(1)2003) and the Sovereign Bases Areas (SBAs) of Cyprus (Ordinance 21/2008), both of which host breeding grounds of the species exclusively in designated areas of the Natura 2000 network of the island (Zaggas et al. 2009; SBAA 2015).

In this study, we aimed to overcome the challenges of assessing the breeding phenology, breeding success, and nest site suitability of sea-cliff nesting raptors using a variety of survey techniques. We studied breeding colonies of Eleonora’s falcon in Cyprus, focusing in particular on the Akrotiri colony, which hosts ~30 pairs (Hadjikyriakou and Kirschel 2016) and comprises our core study site. Specifically, population status was evaluated through surveys for six consecutive years. Nesting habitat selection was also examined with the use of a Digital Surface Model, incorporating topographic factors such as nest elevation, aspect and cliff inclination. In addition, breeding phenology was examined assessing the effects of physical nest characteristics on all the breeding stages. Finally, reproductive success rates were estimated during all breeding stages from territory occupancy to fledging. To accomplish these diverse objectives, we combined boat, ground and aerial surveys with photography and photogrammetry.

## MATERIAL AND METHODS

### Study species

Eleonora’s falcon *(Falco eleonorae* Géné, 1839) is a social raptor with a total population of ~15,000 breeding pairs (Walter 1979; BirdLife International 2017; Touati et al. 2017), breeding primarily on islands and islets in the Aegean Sea (Dimalexis et al. 2008). The breeding population though ranges from Cyprus in the east, westwards across the Mediterranean Sea, the Atlantic coast of Morocco and the Canary Islands (Walter 1979). Colonies are irregularly distributed, ranging from a few pairs up to a few hundred pairs (Vaughan 1961; Walter 1979), typically located on remote and isolated coastal cliffs and islets along migratory routes of small passerines (Vaughan 1961; Walter 1979; Orta 1994). In Cyprus, hosting the easternmost breeding colonies of Eleonora’s falcon, nests are situated on sea cliffs on the south coast of the island, between Cape Gata in the east and Petra tou Romiou in the west (Figure 1).

**Figure 1.**
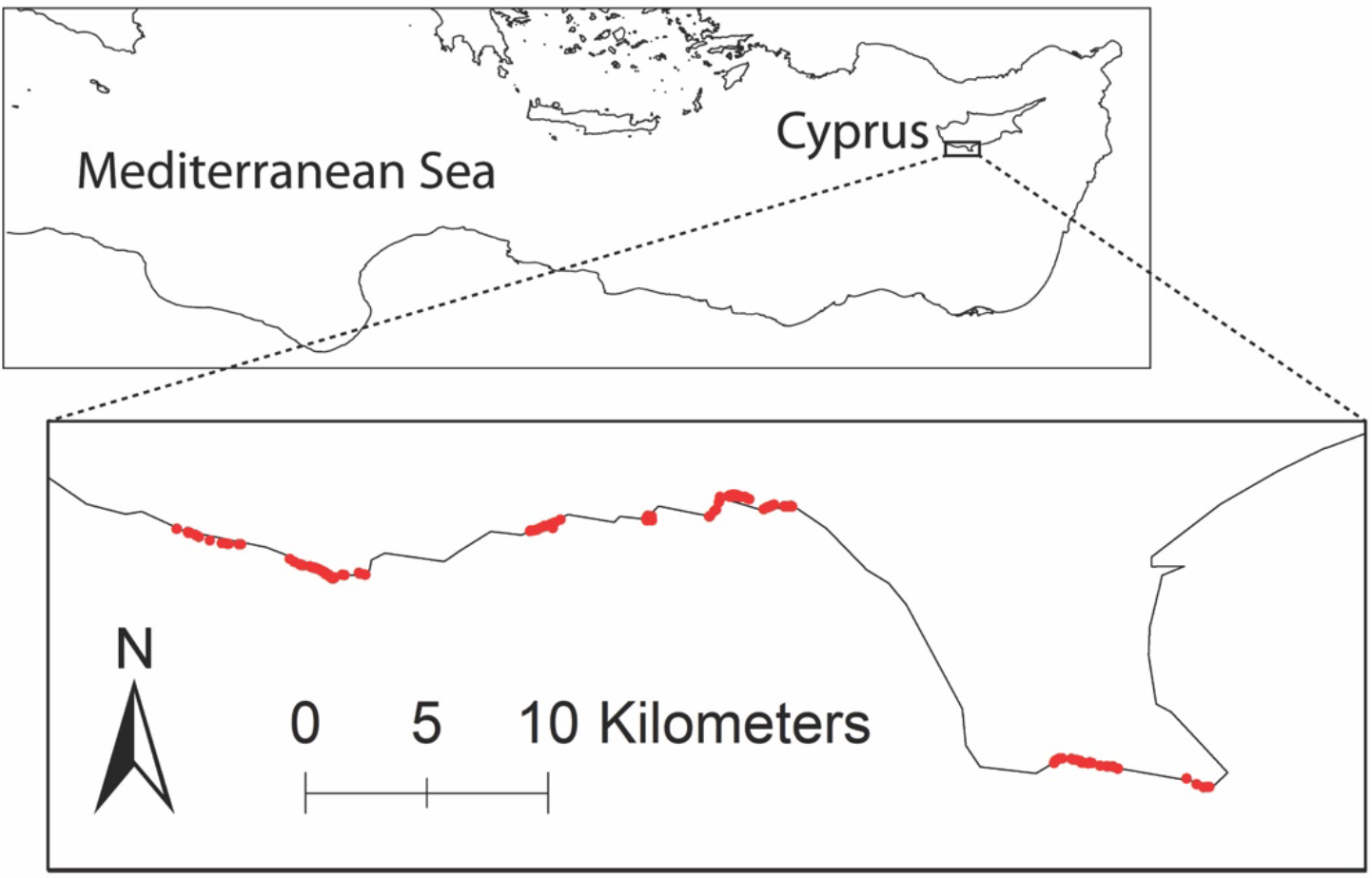
Study area at the southern coast of Cyprus (map boundaries drawn from Eurostat (2015), with nesting sites comprising the three colonies of Cape Aspro, Episkopi and Akrotiri, spread from “Petra tou Romiou” on the west, to “Cape Gata” on the east

Eleonora’s falcon feeds on large (over 10 mm long) insects throughout the year, but gradually switches its diet to small migrating birds, up to about the size of a turtle dove *(Streptopelia turtur),* during the nestling rearing period to cope with the increasing energetic requirements of nestlings (Walter 1979). Although suitable nest sites on coastal cliffs are limited and exposed to winds, breeding pairs prefer to nest there rather than further inland where disturbance and predation are inevitable (Walter 1979; Urios and Martínez-Abraín 2006). Eleonora’s falcons are typically present at their breeding grounds between April and November (Walter 1979), following their migration across continental Africa (Hadjikyriakou et al. 2020a) from their wintering grounds in Madagascar (Kassara et al. 2017; Hadjikyriakou et al. 2020b).

### Population surveys

Population counts were conducted primarily through boat surveys, with the exception of a cliff section with precarious boat access due to shallow waters; this was surveyed from the base of the cliff which was accessible on foot. All population counts were completed during the nestling rearing period, specifically around the last week of September when nestlings were old enough to leave the nest scrape, becoming more visible, but not yet having fledged (Walter 1979). One boat survey per colony in Cyprus (Akrotiri, Episkopi and Cape Aspro) was performed each year for six consecutive breeding seasons (2012 – 2017). Surveys were conducted between 0600 – 1100 hr, counting all adults seen. Our aim was to perform counts from a distance about equal to the cliff height with boat speed 2-4 knots/h (Dimalexis et al. 2008), and stopping to record nestlings.

For population estimates we used survey data from the duration of our study (2012 – 2017), incorporating data from surveys performed previously (2000 – 2011) by other groups using similar methods (Warne 2001; Paton 2003; Pickard and Wilson 2004; Wilson 2005; Miltiadou 2008a; Miltiadou 2008b; Miltiadou 2009; Miltiadou 2011; M. Miltiadou pers. comm.). We used linear trend modelling to analyse the data in TRIM (Pannekoek et al. 2006), with software imputed values for two missing years, 2001 and 2006. Although population counts may give an overview of the condition of a colony, they include non-breeding individuals (Dimalexis et al. 2008), such as sub-adults and floaters (Brown 1974; Postupalsky 1974; Steenhof and Newton 2007). It is the breeding pairs that contribute at any given time to population sustainability (Brown 1974). We thus estimated the number of breeding pairs for all colonies based on the ratio between the number of breeding pairs recorded and the number of adults counted at the more intensively studied Akrotiri colony.

### Nesting habitat selection

Nest site preferences were evaluated using a Digital Surface Model (DSM) of the Akrotiri colony cliffs, created with photogrammetry. DSM in conjunction with GIS is a powerful combination for the prediction of the presence of species (Guisan and Zimmermann 2000). A large body of work has used GIS with remotely sensed environmental data to predict distributions of plants and animals (Saatchi et al. 2008; Kassara et al 2017), and bird behaviour (Smith et al. 2013; Kirschel et al. 2019). Indeed, features of the environment have recently been found to influence Eleonora’s falcon phenology and behaviour during migration and at their wintering grounds (Hadjikyriakou et al. 2020a, Hadjikyriakou et al. 2020b).

The cliffs at Akrotiri stretch for ~10 km from east to west with elevation ranging from sea level to 60 m above sea level. Two sets of low oblique overlapped photographs of the cliff face were taken during two flights with a Cessna plane, using a Canon EOS 40D camera body and a Canon EF70-200mm f/2.8L IS IIUSM lens, from a distance of ~100 m and flight heights of ~100 m and 200 m respectively. Photographs were processed with Agisoft’s Photoscan software using GNSS control points acquired on site (Skarlatos et al. 2013). The final photogrammetric products were an orthophotograph of the study area with a pixel size of 0.5 m and a DSM with 0.5 m. It should be noted that the cliff model created by Photoscan is a full 3D model of the area, but the exported DSM reduces the 3D model to a 2.5D model, losing some information (Mikhail et al. 2001). This is particularly important when the area of interest has steep and vertical slopes (Mikhail et al. 2001). The target cliff area was isolated using the subsequently created contour map, where the top and bottom of the cliffs could be identified and a polygon was selected to limit the Area Of Interest (AOI) to the cliff area only. The DSM was imported into ArcGIS (ESRI 2012) and downsampled to 2 x 2 m for further processing, as this size was considered ecologically meaningful for the birds within the context of this study. Inclination and aspect were calculated within the GIS environment. The DSM was queried to assess suitability of the cliffs using nest positions obtained from six years of surveys. We considered the area of the cliff falling within one standard deviation of the mean for each of the three DSM parameters, i.e., elevation, inclination and aspect as highly suitable.

### Breeding phenology

We investigated the effects of physical nest characteristics and timing of egg laying on breeding success using Generalized Linear Mixed Models (GLMMs) fitted using maximum likelihood estimation via Template Model Builder (GLMMTMB) in R 3.5.1 (R Core Team 2018). Specifically, we tested for the effect of laying date, cliff inclination, nest type, nest aspect and nest elevation, with year as a fixed random factor, on fledgling numbers of successful pairs in GLMMs with a zero-truncated Conway-Maxwell Poisson distribution (Conway and Maxwell 1962). Apart from the physical nest characteristics mentioned above, in the same model we tested for the effect of colony and rainfall during the pre-breeding season. Rainfall is related to insect abundance (Wolda 1978; Wolda 1988), which in turn may affect clutch size (Ristow 2004; Steenhof and Newton 2007). Eleonora’s falcon ranges widely during the pre-breeding period over the southern part of Cyprus feeding on insects (First author personal observations), so we used precipitation data for the southern part of Cyprus for the period from April to August for all the years of this study (2012 – 2017) (Department of Meteorology 2017).

We also built GLMMs in GLMMTMB for the intensively studied colony at Akrotiri for the 2013 breeding season, to test for the effect of laying date, cliff inclination, nest type, nest aspect and nest elevation on clutch size (with zero-truncated Conway-Maxwell Poisson distribution), and on eggs hatched and hatchlings fledged per breeding pair (both with Conway-Maxwell Poisson distribution). In addition, for the Akrotiri colony where we had data for nest usage from 2012-2017, we ran GLMMs in GLMMTMB (zero truncated Conway-Maxwell Poisson distribution) by adding nest occupancy and year as fixed factors, and nest ID as a random factor, to test if nest usage frequency had any effect on number of fledglings per successful pair. Continuous variables in all models i.e. laying date, cliff inclination, nest elevation and rainfall were centered and standardized (Schielzeth 2010). We tested a set of candidate models and model averaging was performed using AICcmodavg 2.2-2 in R on those candidate models with ΔAICc < 4. Significant were those effects of which confidence intervals were not crossing over zero. *P* values were not used because these are only provided for specific models and not by the averaging R package we used.

### Reproductive success rates

Further to the overall population size estimates (total population excluding nestlings), actual breeding pairs (pairs occupying territory) and successful pairs (pairs successfully rearing at least one fledgling) can enhance reproductive success estimates providing the base for the calculation of fledglings per breeding pair, and fledglings per successful pair respectively. For all colonies in Cyprus we recorded active nests identified during the population count surveys between 2012 and 2017 (see above). We recorded nest characteristics and bird related information for each active nest where nestling(s) were spotted, using a protocol modified from HOS (2012). These included weather conditions, nest type (exposed ledge, deep ledge, hole, cave, crevice and under bush), nest aspect (N, NE, E, SE, S, SW, W, NW), cliff inclination (in bands of 10 degrees), visibility within the nest (scale from 1 – 3), remote nest mapping information (GPS location, bearing, distance and angle of observation), as well as nestling and pair presence and activity (Walter 1979; Xirouchakis et al. 2012).

All the nests were photographed for further examination to enhance data accuracy and estimate nestling age (Clark 2007) by comparing the development stage of each nestling with daily video footage from four camera traps installed at Akrotiri colony during the 2013 breeding season. Two of the installed camera traps covered the entire breeding season (see Hadjikyriakou and Kirschel 2016), providing footage for nestlings on a daily basis from hatching until fledging. This way, we were able to estimate nestling age to an accuracy of within three days and consecutively calculate laying, hatching and fledging dates (Green 2004), based on the duration of each breeding stage observed from cameras, as well as on literature (Walter 1968; Walter 1979; Dawson 2004). Because of the location of the nests on steep sea cliffs, using a GPS to directly map nest positions (López-Darias and Rumeu 2010) was not possible. Instead, we calculated nest coordinates and nest elevation using trigonometry after taking distance, angle of view and bearing from the boat using a laser rangefinder and a compass. We note here that at Akrotiri we collected data for the usage frequency of each nest for the whole duration of the study.

In 2013 we studied the breeding stages at Akrotiri from nest occupation to egg laying, hatching and fledging, to identify possible between-stage losses (Postupalsky 1974; Steenhof and Newton 2007; Thorup et al. 2010). Akrotiri is considered the most suitable colony in Cyprus for such systematic work (Flint 1972) because cliffs are relatively low and many nests can be monitored from the cliff top. For each breeding stage we used the following methods (Andersen 2007):

- Territory occupancy was estimated early in the breeding season through cliff-top and boat surveys, recording pairs around nest sites or territorial activity of one partner (Walter 1979; Steenhof and Newton 2007). To increase nest finding efficiency for our core study area, we undertook cliff-top surveys in July and August 2012 along a transect covering the entire length of Akrotiri cliffs recording positions of all adults seen with a handheld GPS.
- Clutch size was recorded for some nests from the cliff top, but for most nests this was not possible. This was not feasible from a boat either, because nests were high up on steep sea cliffs and visibility within the nest from a convenient angle in most cases was not possible (Bibby et al. 2000; Steenhof and Newton 2007). Hence, in addition to cliff-top counts, during the second week of August when most pairs had laid their eggs but had not yet hatched (Wink et al. 1993; Xirouchakis et al. 2012), we used a small custom-made drone (quadcopter) equipped with a Hero 3 GoPro camera. The drone took off and was operated from a boat, approaching nests using wirelessly transmitted live view, obtaining photo and video footage, which was examined later to count the number of eggs in each nest. The highest possible photo and video resolution was used to enhance egg identification (12 MP images and 1440p video resolution at 24 frames per second). Every effort was undertaken to take footage as quickly as possible and to minimise disturbance, with most drone flights lasting less than 5 min, while maintaining a distance to nests of over 10 m (Vas et al. 2015; Radiansyah et al. 2017). Upon drone approach, any parents present vacated the nest and returned a few minutes after its withdrawal following footage capture.
- Counts of hatchlings were undertaken during the first week of September when most nestlings were expected to have hatched (Xirouchakis et al. 2012). These were performed via boat and cliff-top survey using binoculars and telescopes. All nests were photographed to enhance data collection accuracy.
- Number of nestlings fledged was recorded through boat and cliff top-surveys. In Eleonora’s falcon, nestling mortality has been estimated at ~10% in the first 2 weeks, while thereafter mortality is negligible (Ristow 1999). Thereafter nestling numbers are consistent with fledgling numbers (Clark 1974; Ristow 1999; Steenhof and Newton 2007), so the number of nestlings counted from mid-September onwards were considered a reliable estimate of the number of fledglings (Ristow 1999).

## RESULTS

### Population surveys

We counted a mean 232 (*n* = 6, SD = 14.00) adults per season (Figure 2) and found that the population size was stable over the six years (*n* = 6, Wald-test (1) = 1.68, *P* = 0.195). Population size also remained stable when including previous counts back to 2000 *(n* = 16, Wald-test (1) = 1.19, *P* = 0.275). We counted an average of 69 successful nests per year (*n* = 6, Mean = 68.50, SD = 9.77). Because of the challenges of nest detection for this species, the actual total number of breeding pairs is expected to be higher (Walter and Foers 1980). By using Akrotiri as a baseline we estimated that Cyprus hosts 138 breeding pairs (*n* = 6, Mean = 137.8, SD = 8.30).

**Figure 2.**
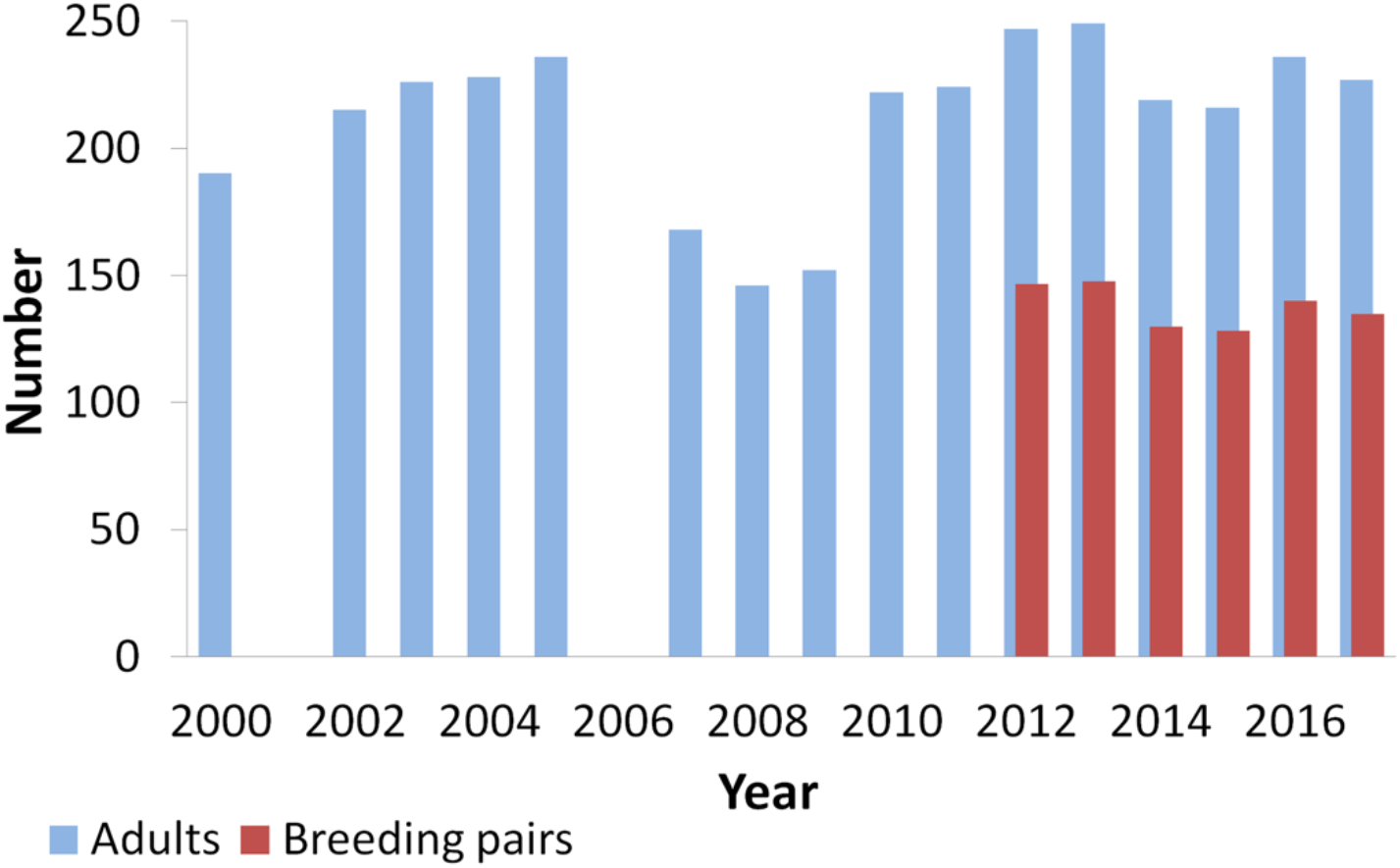
Adults counted during Eleonora’s falcon annual population surveys from yr 2000 onwards, and breeding pairs estimated for the duration of this study (2012 – 2017). Previous counts (2000 – 2011) compiled from surveys performed by other groups (Warne 2001; Paton 2003; Pickard and Wilson 2004; Wilson 2005; Miltiadou 2008a; Miltiadou 2008b; Miltiadou 2009; Miltiadou 2011; M. Miltiadou pers. comm.). Population size was stable over the six years of this study (*n* = 6, Wald-test (1) = 1.68, *P* = 0.195). Population size remained stable also when including previous counts back to 2000 (*n* = 16, Wald-test (1) = 1.19, *P* = 0.275).

### Nesting habitat selection

Almost all Eleonora’s falcon nests in Cyprus were located on near vertical sea cliffs, at elevations ranging from 1 to 100 m, while nest aspect was predominantly southerly (Table 1). However, for the core study area of Akrotiri, most nests were also south-facing but with a slightly more westerly orientation compared to the other two colonies, while the elevation used was on average lower at Akrotiri (Table 1), presumably because the cliffs are lower there than at the other two colonies. Sixty-seven percent of nests were on exposed ledges, while 19.3 % were within holes, 8.4% in deep ledges, 2.6 % in crevices, 1.7 % under bushes and 0.2 % in caves. The DSM on the Akrotiri cliffs indicated that out of the potentially available cliff stretch, just a small fraction (1.14 %) is highly suitable for nesting (Figure 3). This percentage reflects highly suitable cliff sections with regard to elevation, aspect and inclination, but does not provide information on the availability of suitable nest locations (e.g., ledges, holes and crevices), thus actual nest location availability is expected to be even lower.

**Figure 3.**
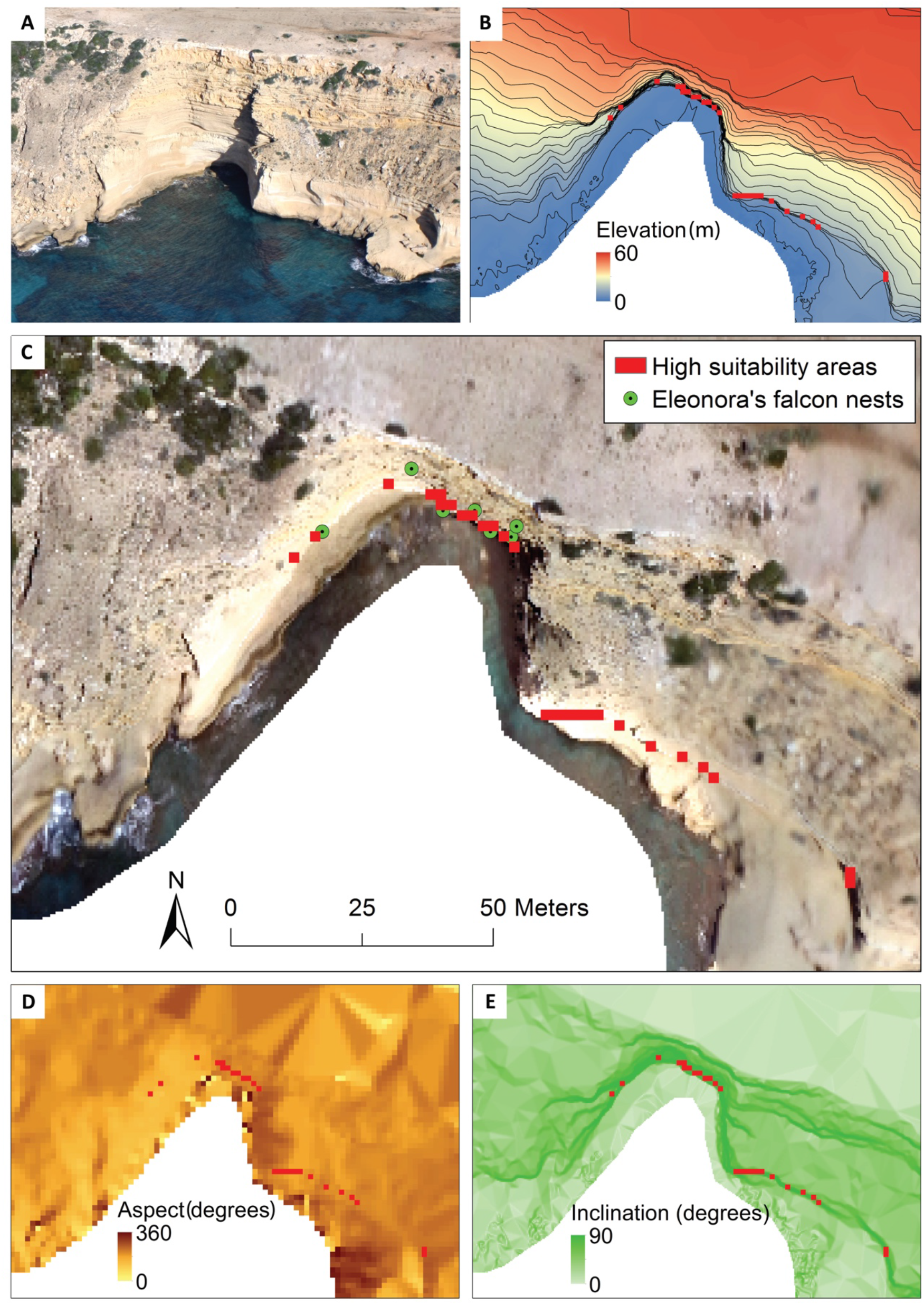
DSM components showing prediction of highly suitable nesting areas of a small section at Akrotiri colony, with A) a photo of the cliff section, B) elevation raster, C) orthophotograph of a section of Akrotiri cliffs incorporating actual nests, D) aspect raster and E) inclination raster. DSM indicated that out of the potentially available cliff stretch, just a small fraction (1.14 %) is highly suitable for nesting.

**Table 1.**
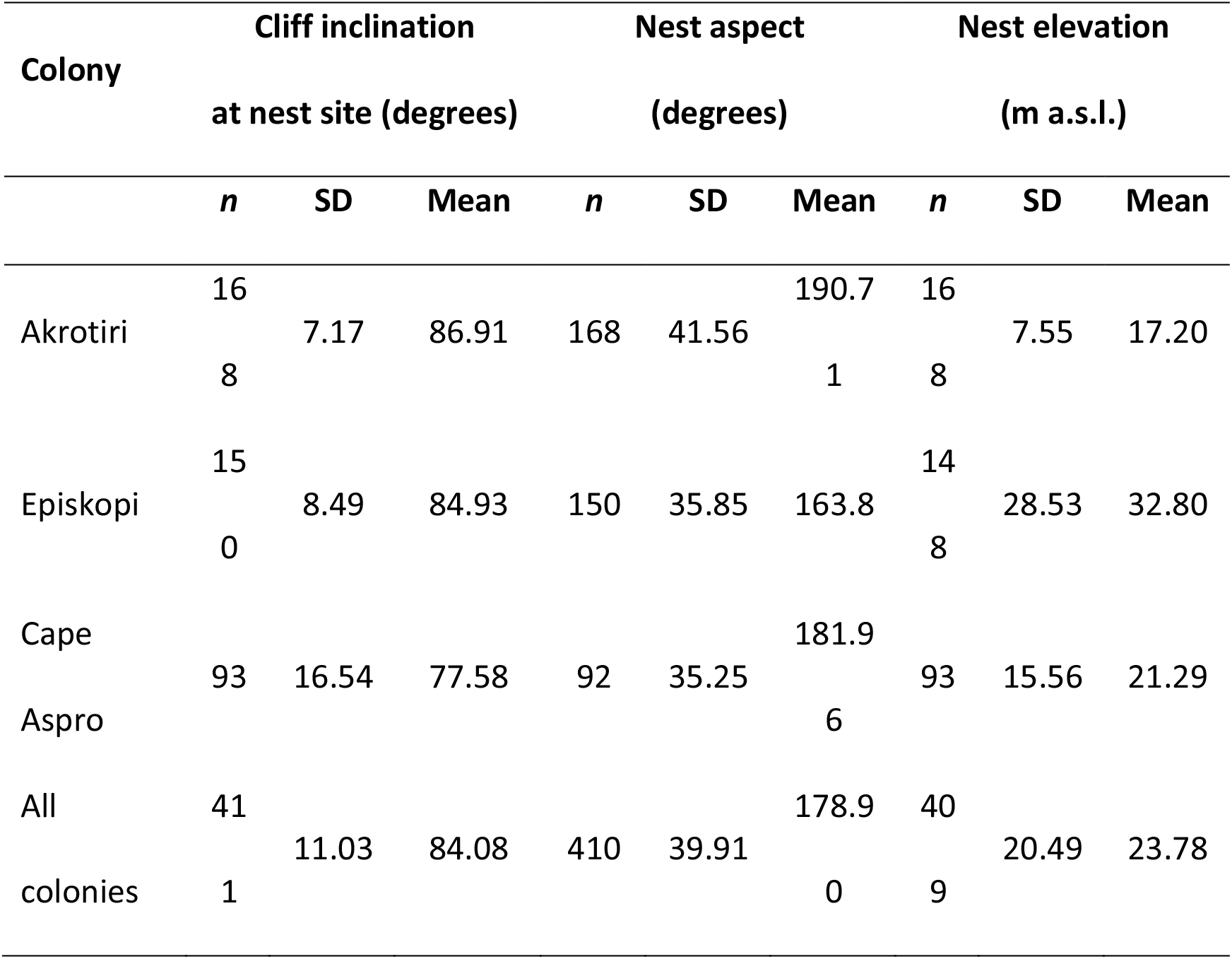
Nest characteristics for colonies in Cyprus (Akrotiri, Episkopi and Cape Aspro) and average for all three colonies

### Breeding phenology

Based on the observed nestling ages for all nesting events between 2012 – 2017 (*n* = 410), eggs hatched between 15^th^ August and 18^th^ September, peaking on 26^th^ August. Given that egg laying typically occurs 30 days earlier (Wink et al. 1993; Orta 1994), the predicted egg-laying peak was on 27^th^ July. Accordingly, fledging normally takes place when nestlings are ~37 day sold (Walter 1979; Orta 1994), therefore, the fledging period is expected to have peaked on 2^nd^ October.

Incorporating the data from all three colonies for the breeding seasons 2012 – 2017, we found a negative correlation of laying date with fledgling numbers for successful pairs, with early breeders more successful, while precipitation during the pre-breeding season was positively correlated with the number of fledglings (Table 2A). One of the nest types (cave) had a significant positive effect on number of fledglings, while Episkopi colony had significantly lower fledgling numbers compared to Akrotiri colony. At Akrotiri, specifically during the 2013 breeding season, we found that cliff inclination and nest elevation were significantly positively associated with clutch size (Table 2B and supplemental material Figure S1), but no factors were significantly associated with hatching success and fledglings per breeding pair (Tables S3 and S4 in supplemental material). We did however find that nesting in cliff holes and the 225 ° nest aspect had a negative effect on fledglings per successful pair at Akrotiri colony over the entire study period (2012 – 2017). Furthermore, there was a significant positive association between number of times a nest was used (nest occupancy) and number of fledglings per successful pair (Figure S2 in supplemental material and Table 2C; full model results in supplemental material, Tables S1 – S5).

**Table 2.** (A) Zero truncated Conway-Maxwell Poisson GLMM significant results for the effect of laying date, cliff inclination, nest type, nest aspect, nest elevation, colony and rainfall during the pre-breeding season on number of fledglings per successful pair for all colonies for breeding seasons 2012 - 2017; (B) Zero truncated Conway-Maxwell Poisson GLMM significant averaged results for the effect of laying date, cliff inclination, nest type, nest aspect and nest elevation on clutch size for Akrotiri colony for 2013 breeding season; (C) Zero truncated Conway-Maxwell Poisson regression model (GLMMTMB) significant averaged results for the effect of laying date, cliff inclination, nest type, nest aspect, nest elevation, year and nest occupancy on number of fledglings per successful pair for Akrotiri colony for breeding seasons 2012 - 2017. Significant effects were those with confidence intervals not crossing over zero (full model results are presented in the supplemental material, Tables S1 - S5)

| Variable | Estimate | SE | Confidence intervals |  |
| --- | --- | --- | --- | --- |
| Number of fledglings per successful pair for all colonies |  |  |  |  |
| (A) | 2012 – 2017 (*n* = 408) |  |  |  |
| Laying date | -0.07 | 0.02 | -0.12 | -0.03 |
| Nest type (cave) | 1.10 | 0.37 | 0.36 | 1.83 |
| Colony (Episkopi) | -0.11 | 0.05 | -0.20 | -0.02 |
| Rainfall | 0.05 | 0.02 | 0.01 | 0.09 |
| Clutch size for Akrotiri colony |  |  |  |  |
| (B) | 2013 (*n* = 27) |  |  |  |
| Cliff inclination | 0.18 | 0.09 | 0.00 | 0.36 |
| Nest elevation | 0.12 | 0.06 | 0.00 | 0.23 |
| Number of fledglings per successful pair for Akrotiri colony |  |  |  |  |
| (C) | 2012 – 2017 (*n* = 165) |  |  |  |
| Nest type (hole) | -0.35 | 0.17 | -0.68 | -0.01 |
| Nest aspect (225 °) | -0.18 | 0.08 | -0.33 | -0.02 |
| Nest occupancy | 0.05 | 0.02 | 0.01 | 0.09 |

### Reproductive success rates

A total of 411 successful nesting events were identified during the study period comprising 729 fledglings. The nests found at Akrotiri colony coincided with areas where adult falcons have been recorded flying during cliff-top surveys early in the breeding season, indicating that adults moving close to the cliff edge can be used to predict the nesting areas. The nests contained 1 – 3 fledglings (*n* = 411, Mean = 1.77, SD = 0.69). During the 2013 breeding season, 35 breeding pairs were identified at Akrotiri, of which 30 were successful in rearing at least one fledgling. The egg stage surveys confirmed that 1 – 3 eggs were laid in each nest (*n* = 27, Mean = 2.44, SD = 0.75). There were on average 1.90 (*n* = 29, SD = 0.86) hatchlings per nest, and 1.80 (*n* = 30, SD = 0.85) fledglings per successful pair. For 2013 we estimated 1.54 (*n* = 35, SD = 0.85) fledglings per breeding pair (Table 3).

**Table 3.** Breeding success results of this study, compared with previous work in Cyprus (Walter and Foers 1980) and the Aegean Sea (Xirouchakis et al. 2012). At the intensively studied Akrotiri colony, where success at all breeding stages was estimated during the 2013 season, results indicate a higher success compared with previous work in the same area and compare favourably with the Aegean Sea. Furthermore, fledglings per successful pair, which was estimated for six consecutive years for all colonies in Cyprus (2012 – 2017), shows similar results.

| Estimator | Area and period of study |  |  |  |
| --- | --- | --- | --- | --- |
|  | Akrotiri | Cyprus (All colonies) | Akrotiri | Aegean Sea |
|  | 2013 | 2012 - 17 | 1977 | 2004 – 07 |
| Clutch size | 2.44 |  |  | 2.43 |
| Eggs hatched per nest | 1.90 |  |  | 1.58 |
| Fledglings per successful pair | 1.80 | 1.77 | 1.53 | 1.84 – 2.00 |
| Fledglings per breeding pair | 1.54 |  | 0.8 – 1.0 | 1.19-1.75 |

## DISCUSSION

Previous research on Eleonora’s falcon in Cyprus had concluded on stable population trends based on adult population numbers (Flint and Stewart 1992; Warne 2001). However, considering the strong site tenacity, low adult mortality and life expectancy of Eleonora’s falcon, a reduction in breeding success can go unnoticed for many years (Ristow et al. 1989). Thus, breeding success estimation is critical for assessing raptor population health. A reduction in breeding success can severely deplete population size and lead to local extinction, as happened with the Peregrine falcon *(Falco peregrinus)* in North America and Western Europe (Brown 1974; del Hoyo et al. 1994; Ambrose et al. 2016). The only previous study that comprehensively assessed the breeding success of Eleonora’s falcon in Cyprus was one based on a survey conducted in 1977 (Walter and Foers 1980). It found that breeding success was lower than the threshold for a sustainable population (Ristow and Wink 1985), raising concerns that this isolated breeding population might be in decline (Gschweng et al. 2011). The projected decline was despite the fact that colonies in Cyprus were considered free of human interference and predation, with nests on steep, isolated sea cliffs (Walter 1979; Barov and Derhé 2011).

In this study we used an integrated approach (Newton et al. 2016) to comprehensively assess the population of Eleonora’s falcon in Cyprus. Our multi-year study found support for population stability, matching historical records from Cyprus going back to 1972 (Corris 1972;Warne 2001), with some expected fluctuation between years (Brown 1974). We found that the temporal breeding pattern of Eleonora’s falcon in Cyprus was almost identical with findings for other colonies, especially in the Aegean Sea (Walter 1968; Walter 1979; Wink and Ristow 2000). We emphasise here the usefulness of indirect calculation of laying, hatching and fledging dates for some 411 successful nests and 729 fledglings, based on photographic and video evidence of nestlings (Hadjikyriakou and Kirschel 2016), minimising impractical intervention and disturbance at each nest (Green 2004). Monitoring the progress of nestling development would not have otherwise been possible, due to either nesting cliff inaccessibility, or lack of resources (Naude et al. 2019).

Clutch size at Akrotiri was at similar levels with colonies in the Aegean Sea (Xirouchakis et al. 2012). By contrast, pairs at the other extreme of the breeding range in the west lay more eggs (Walter 1979). These differences in clutch size can be attributed to the different food resource base. Specifically, bird prey densities are higher and migrant birds are available for longer during the day at colonies in the Canary Islands and the Atlantic coast of Morocco than those in Cyprus (Walter 1979). The migration pulse in Cyprus is restricted to the early morning hours, suggesting that this small hunting window may result in limited hunting opportunities (Walter 1979). In contrast to previous studies (Wink et al. 1985), we did not find an effect of laying date on clutch size, but in agreement with previous studies elsewhere (Walter 1979), we found a significant effect of laying date on fledglings per successful pair, with early breeders raising more fledglings per successful pair. It is possible that laying earlier does not necessarily influence how many eggs a pair lays but does influence offspring survival until they fledge. Cannibalism of young from a later nest performed by a parent feeding her offspring that were laid earlier supports this possibility (Hadjikyriakou and Kirschel 2016).

Hatchling and fledgling numbers per breeding pair at Akrotiri were not affected by any of the variables tested in this study (laying date, cliff inclination, nest type, nest aspect and nest elevation). Nevertheless, hatching success was higher in our study compared to findings in the Aegean Sea, where many pairs lay eggs in exposed nests rather than protected sea cliffs (Walter 1979; Wink and Ristow 2000), and rat predation and sun irradiation are evident (Ristow and Wink 1985). In addition, the number of fledglings per breeding pair at Akrotiri colony was much higher than previous estimates for the same area (Walter and Foers 1980). Thus, our results ought to alleviate concerns raised by previous studies for the possible decline of the Cypriot population (Walter 1979), puzzling researchers since (Ristow and Wink 1985; Gschweng et al. 2011). Furthermore, fledglings per successful pair for all colonies in Cyprus were also higher than previous estimates and on par with estimates for the Aegean Sea colonies (Xirouchakis et al. 2012). Comparing the results from our surveys with those from previous work is not straightforward (Brown 1974), because of methodological differences between studies (Walter 1979; Flint and Stewart 1992; Warne 2001). We believe that the six-year duration of our study, conducted with consistent effort (same standardised methodology, same observers) and combined with an intensive breeding success assessment, provided reliable estimates of breeding success.

Physical nest characteristics had limited effect on breeding success. The positive effect of deeper caves in contrast to the negative effect of smaller, shallower holes, on fledglings per successful pair, indicates that more exposed nests had fewer fledglings, in agreement with studies in other areas (Touati et al. 2017), but our results are from a small sample of nests in caves and holes. The Episkopi colony had fewer fledglings per successful pair, something that needs further investigation possibly in relation to disturbance from human activities which are more intense compared to the other two colonies which are more isolated. Furthermore, in line with similar findings for lesser kestrels (Negro and Hiraldo 1993), nests used more often at Akrotiri had more fledglings. This indicates that experienced adults, which repeatedly occupy the same superior nests year after year (Clark and Peakall 1977), contribute more to the colony’s reproductive success (Perrins 1970). Precipitation during the pre-breeding season also had a positive contribution to the number of fledglings per successful pair, possibly due to its relation to insect abundance (Welch et al. 2017). Clutch size might indeed be affected by insect abundance (Wolda 1978; Wolda 1988), but this becomes more complicated for Eleonora’s falcon, because during the nestling rearing period it switches its diet to migrant birds, meaning a sufficient pulse of migrant birds is needed to raise offspring (Perrins 1970). Thus, clutch size alone cannot predict higher fledgling numbers, because survival depends on avian prey abundance, given the switch in diet from insects to birds.

Our study provides conservation managers with vital information for the protection of Eleonora’s falcon in Cyprus, a designation species of the Special Protection Areas (SPAs) of Akrotiri cliffs, Episkopi cliffs (SBAA 2015) and Cape Aspro – Petra tou Romiou (I.A.CO Ltd and BirdLife Cyprus 2016). The colonies in Cyprus should continue to be monitored, especially in light of new, extensive development projects proposed in the vicinity of the nesting areas (I.A.CO Ltd and BirdLife Cyprus 2016), considerably limiting the major insect feeding grounds identified, such as citrus plantations (First author personal observations). In addition, the observed increasing recreational activity in the sea adjacent to nesting cliffs (I.A.CO Ltd and BirdLife Cyprus 2016), might distress breeding colonies in Cyprus as has been documented for Eleonora’s falcon colonies elsewhere (Martínez-Abraín et al. 2002). It is also worrying that bird poaching levels in the area, including shooting of protected species such as raptors within the wider area around breeding grounds, is one of the highest on the island, and includes the mass persecution of 52 red-footed falcons in 2007 (Miltiadou 2007). From a total of 340 diurnal raptors found injured or dead in Cyprus (Cyprus Veterinary Services unpublished records) over the period of the present study (2012 – 2017), of which 42 % were *Falco* spp., only seven were Eleonora’s falcons, nevertheless authorities need to address these threats. Intraspecific nest predation can be an important factor in breeding success, especially in seasons with limited prey abundance (Gangoso et al. 2015; Hadjikyriakou and Kirschel 2016; Steen et al. 2016; Gangoso and Figuerola 2019), thus the monitoring of the migration pulse at Akrotiri can provide valuable information in understanding population dynamics of the colonies in Cyprus.

In order to assess breeding success of Eleonora’s falcon, we incorporated a combination of survey methods from the ground, boat and air, combined with photography and photogrammetry. This integrated approach helped alleviate the challenges of monitoring breeding success of cliff nesting birds. For example, drones allowed us to more accurately estimate clutch size, and photogrammetric analysis allowed us to identify nest aspect and inclination of inaccessible nesting cliffs. The combination of photogrammetry techniques with fieldwork can inform the assessment of the carrying capacity of a colony and the potential for colony expansion. Thus, DSM use can aid the prediction of breeding area expansion to other potentially suitable areas, with conservation objectives aimed at protecting both current and potentially future breeding sites to maintain population stability. Altogether such methods may help us gain a better understanding of the population trends of cliff nesting species and the threats they may face.

## Notes

### Competing Interest Statement

The authors have declared no competing interest.

### Summary of Updates

Figures included at the end

## REFERENCES

Ambrose, S., C. Florian, R. J. Ritchie, D. Payer, and R. M. O’Brien (2016). Recovery of American Peregrine Falcons along the upper Yukon River, Alaska. The Journal of Wildlife Management 80:609–620.

Andersen, D. E. (2007). Survey techniques. In Raptor Research and Management Techniques (D. M. Bird, K. L. Bildstein, Editors). Hancock House Publishers, Blaine, WA, USA. pp. 89–100.

Barov, B., and M. Derhé (2011). Review of the implementation of species action plans of threatened birds in the European Union (2004–2010). BirdLife International for the European Commission.

Berthold, P. (2001). Bird Migration: A General Survey. Oxford University Press, Oxford, UK.

Bibby, C. J., N. D. Burgess, D. A. Hill, and H. Mustoe (2000). Bird Census Techniques, second edition. Academic Press, London, UK.

Bildstein, K. L. (2006). Migrating Raptors of the World: Their Ecology and Conservation. Cornell University Press, Ithaca, NY, USA.

BirdLife International (2017). European birds of conservation concern: populations, trends and national responsibilities. BirdLife International, Cambridge, UK.

Brown, L. (1974). Data required for effective study of raptor populations. In Proceedings of the Conference on Raptor Conservation Techniques, Fort Collins, Colorado, 22-24 March (Hamerstrom, N. H. Jr., B. E. Harrell, and R. R. Olendorff, Editors). Raptor Research Foundation, Vermillion, SD, USA. pp. 9–20.

Clark, A. (1974). The population and reproduction of the Eleonora’s Falcon in Morocco. Bulletin de la Société des Sciences Naturelles et Physiques du Maroc 54:61–69.

Clark, W. S. (2007). Raptor identification, ageing, and sexing. In Raptor Research and Management Techniques (Bird, D. M., and K. L. Bildstein, Editors). Hancock House Publishers, Blaine, WA, USA. pp. 47–55.

Clark, A. L., and D. B. Peakall (1977). Organochlorine residues in Eleonora’s falcon *Falco eleonorae*, its eggs and its prey. Ibis 119:353–358.

Conway, R. W. and W. L. Maxwell (1962). A queuing model with state dependent service rates. Journal of Industrial Engineering 12:132–136.

Corris, W. F. (1972). Eleonora’s Falcon *Falco eleonorae*. The Cyprus Ornithological Society (1957), Annual Report 1972 19:20–21.

Cramp, S., and K. Simmons, Editors (1980). Handbook of the Birds of Europe, the Middle East and North Africa. The Birds of the Western Palearctic, Volume 2: Hawks to Bustards. Oxford University Press, London, UK.

Danchin, E., and R. H. Wagner (1997). The evolution of coloniality: The emergence of new perspectives. Trends in Ecology and Evolution 12:342–347.

Dawson, A. (2004). Techniques in physiology and genetics. In Bird Ecology and Conservation, a Handbook of Techniques (Sutherland, W. J., I. Newton, and R. E. Green, Editors). Oxford University Press, New York, NY, USA. pp. 211–231.

Department of Meteorology (2017). Monthly average precipitation in Cyprus. Department of MEteorology. Ministry of agriculture, rutal development and environment. Republic of Cyprus. Nicosia, Cyprus.

del Hoyo, J., A. Elliott, and J. Sargatal, Editors (1994). Handbook of the Birds of the World, Volume 2: New World Vultures to Guinea Fowl. Lynx Edicions, Barcelona, Spain.

Dimalexis, A., S. Xirouchakis, D. Portolou, P. Latsoudis, G. Karris, J. Fric, P. Georgiakakis, C. Barboutis, S. Bourdakis, and M. Ivovič (2008). The status of Eleonora’s Falcon (*Falco eleonorae*) in Greece. Journal of Ornithology 149:23–30.

ESRI (2012). ArcGIS, Version 10.1. ESRI Inc, Redlands, CA, USA.

European Parliament (2009). Directive 2009/147/EC of the European Parliament and the Council of 30 November 2009 on the conservation of wild birds. Official Journal of the European Union L20/7-25.

Eurostat (2015). Administrative units. European Commission. http://ec.europa.eu/eurostat/web/gisco/geodata/reference-data/administrative-units-statistical-units/countries#countries14

Flint, P. R. (1972). Second bird report 1971, Appendix 2. The Cyprus Ornithological Society (C.O.S.) 2:120–123.

Flint, P. R., and P. F. Stewart (1992). The Birds of Cyprus: An Annotated Check-list. British Ornithologists’ Union, UK.

Gangoso L, I. Afán, J. M. Grande, and J. Figuerola (2015). Sociospatial structuration of alternative breeding strategies in a color polymorphic raptor. Behavioral Ecology 26:1119–1130.

Gangoso, L., and J. Figuerola (2019). Breeding success but not mate choice is phenotype- and context-dependent in a color polymorphic raptor. Behavioral Ecology. https://doi.org/10.1093/beheco/arz013

Green, R. E. (2004). Breeding biology. In Bird Ecology and Conservation, a Handbook of Techniques (Sutherland, W. J., I. Newton, and R. E. Green, Editors). Oxford University Press, New York, NY, USA. pp. 57–83.

Gschweng, M., F. Tataruch, O. Fröhlich, and E. K. Kalko (2011). Dead embryos despite low contaminant loads in eggs of Eleonora’s Falcon. International Scholarly Research Network Zoology 510202:1–6.

Guisan, A., and N. E. Zimmermann (2000). Predictive habitat distribution models in ecology. Ecological Modelling 135:147–186.

Hadjikyriakou, T. G., and A. N. G. Kirschel (2016). Video evidence confirms cannibalism in Eleonora’s Falcon. Journal of Raptor Research 50:220–223.

Hadjikyriakou, T. G., Nwankwo, E. C., Virani, M. Z. and A. N. G. Kirschel (2020a) Habitat availability influences migration speed, refueling patterns and seasonal flyways of a fly-and-forage migrant. Movement Ecology 8:10.

Hadjikyriakou, T. G., Kassara, C., Rene de Roland, L.-A., Giokas, S., Tsiopelas, N., Evangelidis, A., Thorstrom, R. and A. N. G. Kirschel (2020b). Phenology, variation in habitat use, and daily activity patterns of Eleonora’s falcon overwintering in Madagascar. Landscape Ecology 35:159–172.

HOS (2012). Eleonora’s Falcon recording protocol. Hellenic Ornithological Society, Athens, Greece.

I.A.CO Ltd, BirdLife Cyprus (2016). Management plan: SPA “Akrotirio Aspro - Petra tou Romiou”. Game and Fauna Service, Ministry of Interior, Nicosia, Cyprus.

Kassara, C. A. Dimalexis, J. Fric, G. Karris, C. Barboutis, and S. Sfenthourakis (2011). Nest site preferences of Eleonora’s falcon (Falco eleonorae) on uninhabited islets of the Aegean Sea using GIS and species distribution models. Journal of Ornithology 153:663–675.

Kassara, C., L. Gangoso, U. Mellone, G. Piasevoli, T. G. Hadjikyriakou, N. Tsiopelas, S. Giokas, P. López-López, V. Urios, J. Figuerola, R. Silva, et al. (2017). Current and future suitability of wintering grounds for a long-distance migratory raptor. Scientific Reports 7:8798.

Kirschel, A.N.G., N, Seddon, and J. A. Tobias (2019). Range-wide spatial mapping reveals convergent character displacement of bird song. Proceedings of the Royal Society B: Biological Sciences 286: 20190443.

López-Darias, M., and B. Rumeu (2010). Status and population trend of Eleonora’s Falcon *Falco eleonorae* in the Canary Islands. Ornis Fennica 87:35–40.

Martínez-Abraín, A., V. Ferris, and R. Belenguer (2002). Is growing tourist activity affecting the distribution or number of breeding pairs in a small colony of the Eleonora’s Falcon? Animal Biodiversity and Conservation 25:47–51.

Mikhail, E. M., J. S. Bethel, and J. C. McGlone (2001). Introduction to Modern Photogrammetry. John Wiley and Sons Inc, New York, NY, USA.

Miltiadou, M. (2007). Red-footed Falcon (*Falco vespertinus)* slaughter at Phassouri. Cyprus BirdLife. BirdLife Cyprus, Nicosia, Cyprus. pp. 16–17.

Miltiadou, M. (2008a). Eleonora’s Falcon *falco eleonorae*. Cyprus Bird Report 2007. BirdLife Cyprus, Nicosia, Cyprus. pp. 53–54.

Miltiadou, M. (2008b). The Eleonora’s Falcon *Falco eleonorae* breeding count 29th August 2008. Monthly Newsletter September 2008. BirdLife Cyprus, Nicosia, Cyprus. pp. 15–16.

Miltiadou, M. (2009). The Eleonora’s Falcon *Falco eleonorae* breeding count 28 August 2009. Cyprus BirdLife. BirdLife Cyprus, Nicosia, Cyprus. p. 11.

Miltiadou, M. (2011). Autumn 2010 summary results of Akrotiri migratory raptor census and Eleonora’s Falcon breeding census 2010. Cyprus BirdLife. BirdLife Cyprus, Nicosia, Cyprus. p. 8.

Naude, V. N., L. K. Smyth, E. A. Weideman, B. A. Krochuk, and A. Amar (2019). Using web- sourced photography to explore the diet of a declining African raptor, the Martial Eagle *(Polemaetus bellicosus)*. The Condor: Ornithological Applications 121:1–9.

Negro, J., and F. Hiraldo (1993). Nest-site selection and breeding success in the Lesser Kestrel *Falco naumanni*. Bird Study 40:115–119.

Newton, I. (2004). Population limitation in migrants. Ibis 146:197–226.

Newton, I. (2010). Population Ecology of Raptors. Bloomsbury Publishing, London, UK

Newton, I., M. J. McGrady, and M. K. Oli (2016). A review of survival estimates for raptors and owls. Ibis 158:227–248.

Orta, J. (1994). Eleonora’s falcon *Falco* eleonorae. In Handbook of the Birds of the World, Volume 2: New World Vultures to Guinea Fowl (del Hoyo, J., A. Elliott, and J. Sargatal, Editors). Lynx Edicions, Barcelona, Spain. p. 266.

Pannekoek, J., A. J. van Strien, and A. W. Gmelig Mayling (2006). TRends and Indices for Monitoring data, Version 3.54. Statistics Netherlands.

Paton, A. S. (2003). Survey of the Eleonora’s falcon breeding sites in Cyprus - 2002. Royal Airforce Ornothological Society Newsletter 75:4–7.

Perrins, C. (1970). The timing of birds’ breeding seasons. Ibis 112:242–245.

Pickard, A. M., and J. Wilson (2004). The 2003 survey of Eleonora’s Falcon breeding sites in Cyprus. The Osprey 4:39–40.

Postupalsky, S. (1974). Raptor reproductive success: Some problems with methods, criteria and terminology. In Proceedings of the Conference on Raptor Conservation Techniques, Fort Collins, Colorado, 22-24 March (Hamerstrom, N. H. Jr., B. E. Harrell, and R. R. Olendorff, Editors). Raptor Research Foundation, Vermillion, SD, USA. pp. 21–31.

R Core Team (2018). R: a language and environment for statistical computing, Version 3.5.1. R Foundation for Statistical Computing, Austria.

Radiansyah, S., M. D. Kusrini, and L. B. Prasetyo (2017). Quadcopter applications for wildlife monitoring. IOP Conference Series: Earth and environmental science 54 012066:1–8.

Ristow, D. (1999). International species action plan, Eleonora’s Falcon *(Falco eleonorae)*. Birdlife International, Council of Europe, Cambridge, UK.

Ristow, D. (2004). On the insect diet of Eleonora’s Falcon *Falco eleonorae* and its importance for coloniality. In Raptors Worldwide (Chancellor, R. D. and B. U. Mayburg, Editors). The World Working Group on Birds of Prey / MME, Berlin, Germany. pp. 705–712.

Ristow, D., W. Scharlau, and M. Wink (1989). Population structure and mortality of Eleonora’s Falcon *Falco eleonorae*. In Raptors in the Modern World, (Mayburg, B. U., and R.D. Chancellor, Editors). The World Working Group on Birds of Prey, Berlin, Germany. pp. 321–326.

Ristow, D., and M. Wink (1985). Breeding success and conservation management of Eleonora’s Falcon. International Council for Bird Preservation technical publication 5:147–152.

Ristow, D., M. Wink, C. Wink, and H. Friemann (1983). Biologie des Eleonorenfalken *(Falco eleonorae):* 14. das brutreifealter der weibchen. Journal Für Ornithologie 124:291–293.

Saatchi, S., W. Buermann, H. ter Steege, S. Mori, and T. B. Smith (2008). Modeling distribution of Amazonian tree species and diversity using remote sensing measurements. Remote Sensing of Environment 112:2000–2017.

Sanderson, F. J., P. F. Donald, D. J. Pain, I. J. Burfield, and F. P. Van Bommel (2006). Long-term population declines in afro-palearctic migrant birds. Biological Conservation 131:93–105.

SBAA (2015). Protection and management of nature and wildlife (special areas of conservation) order 26 of 2015. Sovereign Bases Areas Administration, Episkopi, Cyprus.

Schielzeth H. Simple means to improve the interpretability of regression coefficients. Methods Ecol Evol 2010;1:103–13.

Skarlatos, D., E. Procopiou, G. Stavrou, and M. Gregoriou (2013). Accuracy assessment of minimum control points for UAV photography and georeferencing. PROC SPIE 8795, first international conference on remote sensing and geoinformation of the environment (RSCy2013), 879514, 5 August 2013. http://dx.doi.org/10.1117/12.2028988

Smith, T. B., R. J. Harrigan, A. N. G. Kirschel, W. Buermann, S. Saatchi, D. T. Blumstein, S. R. de Kort, and H. Slabbekoorn (2013). Predicting bird song from space. Evolutionary Applications 6:865–874.

StataCorp (2013). Stata, Version. 13. College Station, TX, USA.

Steen, R., A. Miliou, T. Tsimpidis, V. Sel ås and G. A. Sonerud (2016). Non-parental infanticide in colonial Eleonora’s Falcons *(Falco eleonorae)*. Journal of Raptor Research 50:217–220.

Steenhof, K., and I. Newton (2007). Assessing nesting success and productivity. In Raptor Research and Management Techniques (Bird, D. M., and K. L. Bildstein, Editors). Hancock House Publishers, Blaine, WA, USA. pp. 181–192.

Swatschek, I., D. Ristow, W. Scharlau, C. Wink, and M. Wink (1993). Populationsgenetik und vaterschaftsanalyse beim Eleonorenfalken *(Falco eleonorae)*. Journal Für Ornithologie 134:137–143.

Thorup, K., P. Sunde, L. B. Jacobsen, and C. Rahbek (2010). Breeding season food limitation drives population decline of the Little Owl *Athene noctua* in Denmark. Ibis 152:803–814.

Touati, L., R. Nedjah, F. Samraoui, A. H. Alfarhan, L. Gangoso, J. Figuerola, and B. Samraoui (2017). On the brink: Status and breeding ecology of Eleonora’s Falcon *Falco eleonorae* in Algeria. Bird Conservation International 27:594–606.

Urios, G., and A. Martínez-Abraín (2006). The study of nest-site preferences in Eleonora’s Falcon *Falco eleonorae* through digital terrain models on a western mediterranean island. Journal of Ornithology 147:13–23.

Vas, E., A. Lescroel, O. Duriez, G. Boguszewski, and D. Gremillet (2015). Approaching birds with drones: First experiments and ethical guidelines. Biology Letters 11 20140754:1–4.

Vaughan, R. (1961). Falco eleonorae. Ibis 103:114–128.

Walter, H. (1968). Zur abhängigkeit des Eleonorenfalken *(Falco eleonorae)* vom Mediterranen vogelzug. Journal Für Ornithologie 109:323–365.

Walter, H. (1979). Eleonora’s Falcon: Adaptations to Prey and Habitat in a Social Raptor. University of Chicago Press, Chicago, IL, USA.

Walter, H., and R. Foers (1980). *Falco eleonorae* on Cyprus: population size and breeding success. The Royal Air Force Ornithological Society Journal 11:88–95.

Warne, A. P. (2001). A survey of Eleonora’s Falcon breeding sites. Annual Report 2000. Cyprus Ornithological Society (1957), Nicosia, Cyprus. pp. 114–115.

Welch, B., C. W. Boal, and B. R. Skipper (2017). Environmental influences on the nesting phenology and productivity of Mississipi Lites *(Ictinia mississippiensis)*. The Condor: Ornithological Applications 119:298–307.

Wilson, J. (2005). Conservationists count Eleonora’s Falcons. BirdLife Cyprus News Autumn 2005. BirdLife Cyprus, Nicosia, Cyprus. p. 13.

Wink, M., H. Biebach, F. Feldmann, W. Scharlau, I. Swatschek, C. Wink, and D. Ristow (1993). Contribution to the breeding biology of Eleonora’s Falcon *(Falco eleonorae)*. In Proceedings of the Conference on Biology and Conservation of Small Falcons, 6-8 September 1991 (Nicholls, M. K. and R. Clarke, Editors). The Hawk and Owl Trust, London, UK. pp. 59–72.

Wink, M., and D. Ristow (2000). Biology and molecular genetics of Eleonora’s Falcon *Falco eleonorae,* a colonial raptor of Mediterranean islands. In Raptors at Risk (Chancellor R. D., and B. U. Mayburg, Editors). The World Working Group on Birds of Prey / Hancock House. pp. 653–668.

Wink, M., D. Ristow, and W. Scharlau (1987). Population structure in a colony of Eleonora’s Falcon *(Falco eleonorae)*. Suppl Ric Biol Selvaggina 12:301–306.

Wink, M., D. Ristow, and C. Wink (1985). Biology of Eleonora’s Falcon *(Falco eleonorae).* Variability of clutch size, egg dimensions and egg coloring. Raptor Research 19:8–14.

Wink, M., C. Wink, and D. Ristow (1980). Biologie des Eleonorenfalken *(Falco eleonorae).* Die Gelegegröße in relation zum nahrungsangebot, jagderfolg und gewicht der altfalken. Journal of Ornithology 121:387–390.

Wolda, H. (1978). Seasonal fluctuations in rainfall, food and abundance of tropical insects. Journal of Animal Ecology 47:369–381.

Wolda, H. (1988). Insect seasonality: Why? Annual Review of Ecology and Systematics 19:1–18.

Xirouchakis, S. (2004). Causes of raptor mortality in Crete. In Raptors Worldwide (Chancellor, R. D., and B. U. Meyburg, Editors) The World Working Group on Birds of Prey / MME, Berlin, Germany. pp. 1–13.

Xirouchakis, S., J. Fric, C. Kassara, D. Portolou, A. Dimalexis, G. Karris, C. Barboutis, P. Latsoudis, S. Bourdakis, and E. Kakalis (2012). Variation in breeding parameters of Eleonora’s Falcon *(Falco eleonorae)* and factors affecting its reproductive performance. Ecological Research 27:407–416.

Zaggas, T., P. Ganatsas, T. Tsitsoni, C. Vlachos, D. Gioulatos, A. Mantzavelas, E. Lazaridou, M. Iacovou, A. Kramvias, N. Peristianis, E. Eftychiou, and G. Faidonos (2009). Management plan for the area Akrotirio Apro - Petra tou Romiou (CY500005). Ministry of Agriculture, Natural Resources and Environment, Environment Service, Republic of Cyprus, Thesalloniki, Greece.

