## Supplementary material for "Breeding success of Eleonora’s falcon (*Falco eleonorae*) in Cyprus revisited using survey techniques for cliff-nesting species" (from another version of this preprint: version 1, 2020-05-05)

**Supplemental material**

Table S1. Zero truncated Poisson regression model (GLMMTMB) averaged results (for models with ΔAICc < 4) for the effect of laying date, cliff inclination, nest type, nest aspect, nest elevation, colony and rainfall during the pre-breeding season on number of fledglings per successful pair for all colonies in Cyprus for breeding seasons 2012 – 2017. Significant effects (confidence intervals not crossing over zero) are presented with bold font

| **Number of fledglings per successful pair for all colonies (2012 – 2017) (*n* = 408)** | | | | |
|---|---|---|---|---|
| **Variable** | **Estimate** | **SE** | **Confidence intervals** | |
| Intercept | 0.59 | 0.08 | 0.44 | 0.74 |
| **Laying date** | **-0.07** | **0.02** | **-0.12** | **-0.03** |
| Cliff inclination | 0.04 | 0.03 | -0.01 | 0.09 |
| Nest type (crevice) | 0.20 | 0.21 | -0.20 | 0.61 |
| Nest type (deep ledge) | 0.09 | 0.19 | -0.28 | 0.46 |
| Nest type (exposed ledge) | 0.09 | 0.17 | -0.25 | 0.43 |
| **Nest type (cave)** | **1.10** | **0.37** | **0.36** | **1.83** |
| Nest type (hole) | 0.06 | 0.18 | -0.29 | 0.41 |
| Nest elevation | 0.00 | 0.02 | -0.04 | 0.04 |
| **Colony (Episkopi)** | **-0.11** | **0.05** | **-0.20** | **-0.02** |
| Colony (Cape Apsro) | -0.06 | 0.06 | -0.17 | 0.05 |
| **Rainfall** | **0.05** | **0.02** | **0.01** | **0.09** |

Table S2. Zero truncated Poisson regression model (GLMMTMB) averaged results (for models with ΔAICc < 4) for the effect of laying date, cliff inclination, nest type, nest aspect and nest elevation on clutch size for Akrotiri colony for 2013 breeding season. Significant effects (confidence intervals not crossing over zero) are presented with bold font

| **Clutch size for Akrotiri colony (2013) (*n* = 27)** | | | | |
|---|---|---|---|---|
| **Variable** | **Estimate** | **SE** | **Confidence intervals** | |
| Intercept | 0.91 | 0.05 | 0.81 | 1.01 |
| Laying date | -0.01 | 0.05 | -0.12 | 0.09 |
| **Cliff inclination** | **0.18** | **0.09** | **0.00** | **0.36** |
| **Nest elevation** | **0.12** | **0.06** | **0.00** | **0.23** |

Table S3. Poisson regression model (GLMMTMB) averaged results (for models with ΔAICc < 4) for the effect of laying date, cliff inclination, nest type, nest aspect and nest elevation on number of eggs hatched for Akrotiri colony for 2013 breeding season. No significant effect found in this dataset

| **Number of eggs hatched for Akrotiri colony (2013) (*n* =27)** | | | | |
|---|---|---|---|---|
| **Variable** | **Estimate** | **SE** | **Confidence intervals** | |
| Intercept | 0.72 | 0.07 | 0.59 | 0.85 |
| Laying date | -0.04 | 0.07 | -0.18 | 0.09 |
| Cliff inclination | 0.12 | 0.11 | -0.09 | 0.33 |
| Nest type (deep ledge) | 0.21 | 0.17 | -0.12 | 0.54 |
| Nest type (hole) | -0.17 | 0.17 | -0.50 | 0.15 |
| Nest elevation | 0.07 | 0.08 | -0.08 | 0.23 |

Table S4. Poisson regression model (GLMMTMB) averaged results (for models with ΔAICc < 4) for the effect of laying date, cliff inclination, nest type, nest aspect and nest elevation, on number of fledglings per breeding pair for Akrotiri colony for 2013 breeding season. No significant effect found in this dataset

| **Number of fledglings per breeding pair for Akrotiri colony (2013) (*n* = 31)** | | | | |
|---|---|---|---|---|
| **Variable** | **Estimate** | **SE** | **Confidence intervals** | |
| Intercept | 0.59 | 0.06 | 0.46 | 0.71 |
| Laying date | -0.10 | 0.07 | -0.24 | 0.03 |
| Cliff inclination | 0.03 | 0.09 | -0.15 | 0.21 |
| Nest type (deep ledge) | 0.15 | 0.14 | -0.13 | 0.43 |
| Nest type (hole) | -0.20 | 0.14 | -0.48 | 0.07 |
| Nest elevation | 0.13 | 0.07 | -0.01 | 0.27 |

Table S5. Zero truncated Poisson regression model (GLMMTMB) averaged results (for models with ΔAICc < 4) for the effect of laying date, cliff inclination, nest type, nest aspect, nest elevation, year and nest occupancy on number of fledglings per successful pair for Akrotiri colony for breeding seasons 2012 – 2017. Significant effects (confidence intervals not crossing over zero) are presented with bold font

| **Number of fledglings per successful pair for Akrotiri colony (2012 – 2017) (*n* = 165)** | | | | |
|---|---|---|---|---|
| **Variable** | **Estimate** | **SE** | **Confidence intervals** | |
| Intercept | 0.55 | 0.19 | 0.17 | 0.93 |
| Laying date | -0.05 | 0.03 | -0.11 | 0.01 |
| Cliff inclination | 0.05 | 0.06 | -0.07 | 0.18 |
| Nest type (deep ledge) | -0.14 | 0.19 | -0.52 | 0.23 |
| Nest type (exposed ledge) | -0.29 | 0.16 | -0.61 | 0.04 |
| **Nest type (hole)** | **-0.35** | **0.17** | **-0.68** | **-0.01** |
| Nest aspect (180°) | -0.07 | 0.08 | -0.22 | 0.09 |
| **Nest aspect (225°)** | **-0.18** | **0.08** | **-0.33** | **-0.02** |
| Nest aspect (270°) | -0.19 | 0.13 | -0.44 | 0.06 |
| Nest elevation | - 0.10 | 0.09 | -0.27 | 0.07 |
| Year | -0.02 | 0.02 | -0.05 | 0.02 |
| **Nest occupancy** | **0.05** | **0.02** | **0.01** | **0.09** |

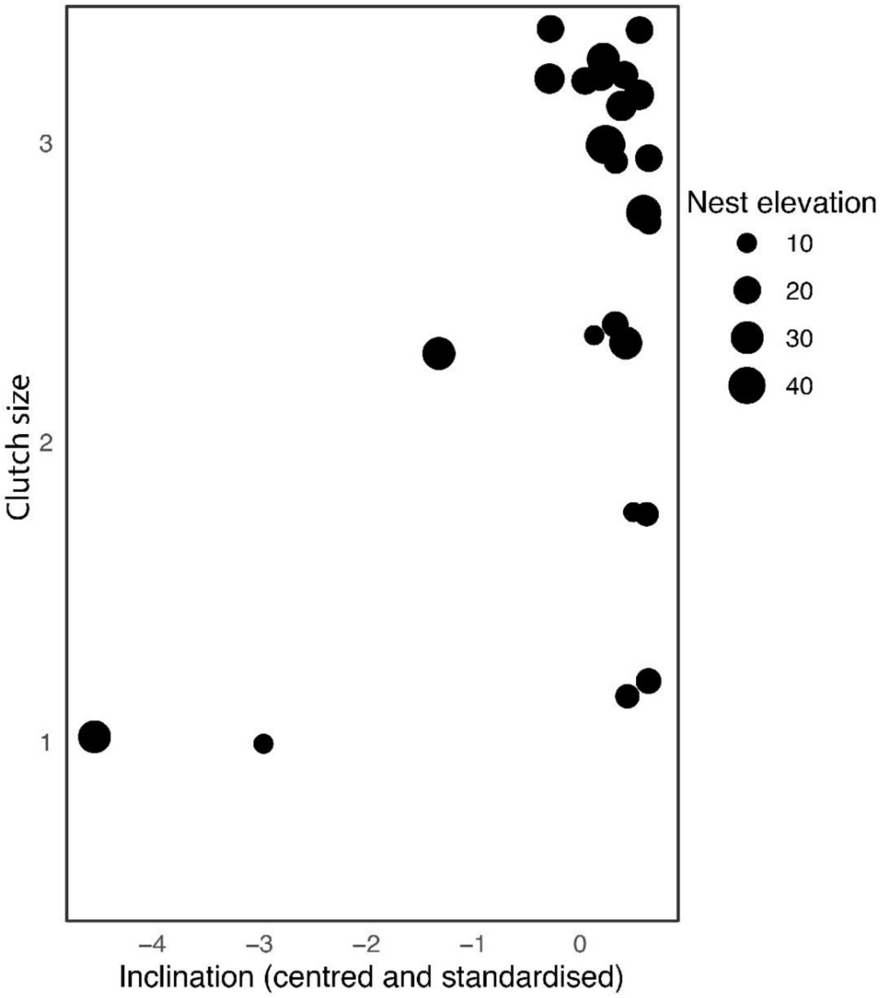


Figure S1. Scatter plot for the effect of cliff inclination and nest elevation on clutch size for Akrotiri colony for 2013 breeding season (The plot uses jitter to visualise points that would otherwise not be visible). Clutch size is positively correlated both with cliff inclination and nest elevation (m a.s.l.).

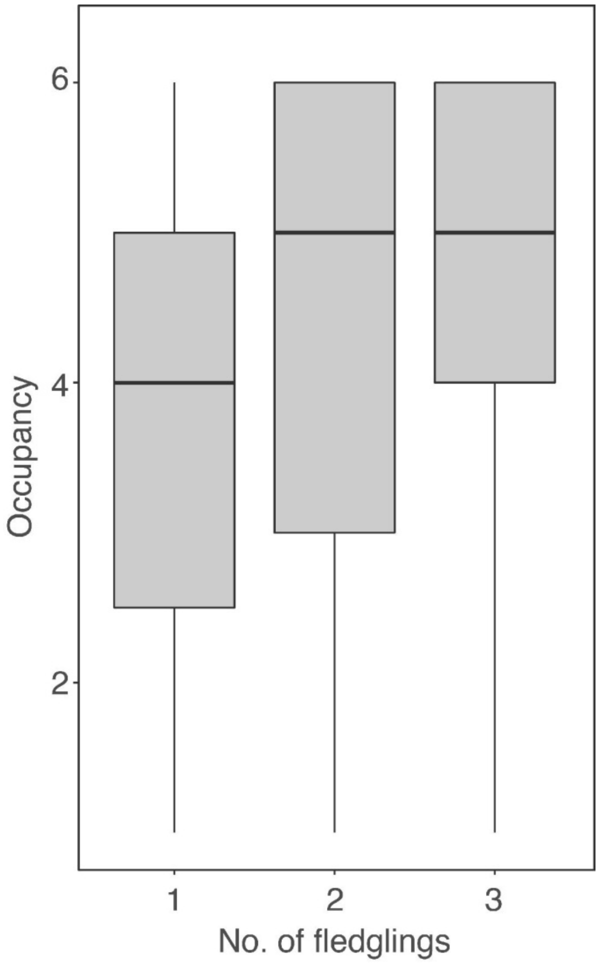


Figure S2. Boxplot for the effect of nest occupancy on number of fledglings per successful pair for Akrotiri colony over the entire study period (2012 – 2017). Number of fledglings is positively correlated with the number of times a nest was used.
